# Notch canonical activity in a subset of glial cells regulates short-term memory in *Drosophila*

**DOI:** 10.1101/2025.10.10.681599

**Authors:** Andrea Aguirre, Dylan Girard, Sandrine Parrot, Claire-Angeline Clugnet, Xavier Biolchini, Hui Li, Laurent Seugnet

## Abstract

Notch is a transmembrane receptor expressed at the cell surface that mediates transcriptional responses upon binding to its ligands, in a variety of contexts. Evolutionarily conserved, Notch plays a key role in numerous cell fate decisions during development and is also required in post-mitotic brain neurons for the consolidation of long-term memory and long-term habituation. Notch signaling is highly expressed in glial cells, where it plays a key role in regulating their development and proliferation. More recently, a Notch-dependent neuroglial pathway has been implicated in modulating the susceptibility of short-term memory to sleep deprivation in *Drosophila*. In this study, we demonstrate that canonical Notch signaling in glia— mediated by the Delta ligand and the transcription factor Suppressor of Hairless—is activated in a subset of cortex and ensheathing glial cells. This signaling is essential for the formation of short-term memory in the aversive phototaxic suppression assay, which is sensitive to sleep deprivation. Dopaminergic transmission is known to be required for this type of learning and is negatively impacted by sleep deprivation, suggesting a possible interaction between Notch and dopamine pathways. Supporting this idea, we find that modulating dopaminergic transmission downregulates canonical Notch activity in glial cells. Conversely, activating Notch signaling in glia near dopaminergic neurons prevents the learning impairments typically caused by sleep deprivation. Notably, Notch signaling itself does not appear to alter dopamine levels in the brain. Together, these findings indicate that canonical Notch signaling in a specific subset of glial cells is essential for short-term memory formation and is modulated by dopaminergic signaling. This suggests that sleep loss–induced disruption of dopaminergic transmission impairs learning by downregulating canonical Notch signaling. Since *Notch* homologs are highly expressed in mammalian glia, this pathway may be conserved and functionally relevant in other species, including humans.

## Introduction

Notch receptors are large single trans-membrane domain proteins containing EGF-like motifs in the extracellular domain, Ankyrin repeats in the intracellular domain as well as several other evolutionary conserved motifs. The nature of the signaling cascade downstream of the Notch receptor depends on the cellular context. The canonical pathway requires the binding of the Notch receptors to the Delta-Serrate-Lag2 (DSL) family of trans-membrane ligands (such as Delta or Serrate in Drosophila), which triggers secretase-mediated cleavages leading to the shedding of the extracellular domain and the release of the Notch intracellular domain into the cytoplasm. The Notch intracellular domain then translocates to the nucleus where it associates with the CBF1/Suppressor of Hairless/LAG-1 complex, including the transcription factor suppressor of hairless (Su(H), RBP-Jκ in mammals) and activates the transcription of targets genes such as genes of the HES family (*hairy* and *enhancer of split*). Several non-canonical Notch pathways have also been described, so-called because they do not involve the conventional DSL ligands, the cleavage of the receptor, or the CSL complex, and/or involves crosstalks with other signaling pathway such as Wnt^1^.

Notch signaling has been shown to play multiple roles in developing tissues and undifferentiated cells where it has been intensively studied. Since more than a decade and in multiple independent studies, Notch signaling has also been shown to be required in the adult brain, where it plays a critical role in the physiology of learning and memory, both in Drosophila and in the mouse^2^. In *Drosophila*, Notch signaling has been shown to be specifically required for the consolidation of long term memories following associative olfactory conditioning or courtship conditioning, while being dispensable for both short term and anesthesia resistant memory, another type of consolidated memory that does not require protein synthesis^3,4^. Notch is also required for the olfactory glomeruli morphological and physiological plasticity following long time exposure to specific odors^5,6^. Neuronal plasticity relies on sleep and interestingly, reports in *Drosophila*^7^ and in *C. elegans*^8^ have provided evidence that Notch is involved in sleep regulation as well. In particular, we have shown that Notch signaling modulates the sleep loss vulnerability of a learning paradigm requiring flies to inhibit an instinctive attraction toward light: Aversive Phototaxic Suppression (APS)^7^.

Importantly most of the studies so far have used tools with no or limited cell specificity^3,9–11^, thus limiting the ability to interpret the results at the cellular level. In the mouse, Notch receptors and members of the signaling cascade are expressed at high level in glia^12^, while the ligands are predominantly expressed in neurons, similar to what has been observed in *Drosophila* and suggesting that Notch could be mediating neuron-glia interactions^7^. In this report we show that Notch canonical signaling is activated in a subset of glial cells, is dependent on the ligand Delta, and that APS depends on the activation of this pathway in cortex and ensheathing glia. We also obtained evidence that dopaminergic transmission is modulating Notch signaling in glia.

## Materials and methods

### Fly stocks and husbandry

Flies were reared at 25°C, 50 – 60% humidity, in a 12 h light/dark cycle in a standard food containing inactivated yeast, molasses, sucrose, and agar. Fly stocks: we used the following fly stocks from the Bloomington Drosophila Stock Center: P{w[+m*]=NRE-EGFP.S}5A (NRE-GFP, RRID:BDSC_30727); P{w[+m*]=NRE-EGFP.S}1 (NRE-GFP, RRID:BDSC_30728); P{UAS-N.LV}2, P{lexAop-hrGFP}2c; P{lexAop-hrGFP}3 (PRRID:BDSC_29965); P{w[+m*]=GAL4}repo (repo-Gal4, (RRID:BDSC_7415);P{y[+t7.7] w[+mC]=GMR75H11-GAL4}attP2 (Eaat1-Gal4, RRID:BDSC_39914); P{TRiP.HMS00009}attP2 (RRID:BDSC_33616, UAS-N-RNAi (1)); P{UAS-N.RNAi.P}14E (RRID:BDSC_7078, UAS-N-RNAi (2));; P{tubP-GAL80[ts]} (Tub-Gal80^ts^, RRID:BDSC_7016); P{y[+t7.7] v[+t1.8]=TRiP.HM05110}attP2 (RRID:BDSC_28900, UAS-Su(H)-RNAi (1)); *Delta*^*RF*^ (RRID:BDSC_5603); P{y[+t7.7] v[+t1.8]=TRiP.JF02867}attP2 (UAS-Dl-RNAi, RRID:BDSC_28032); P{y[+t7.7] v[+t1.8]=TRiP.GL00520}attP40 (UAS-Dl-RNAi, RRID:BDSC_36784); P{UAS-Delta.J} (UAS-Delta, RRID:BDSC_26694); from the Vienna Drosophila Ressource Center : UAS-Su(H)-RNAi^KK101550^ (UAS-Su(H)-RNAi (2)); from the Kyoto Drosophila Stock Center : P{w[+mW.hs]=GawB}NP2222 (NP2222); P{w[+mW.hs]=GawB}NP6520 (NP6520). We thank the colleagues who provided the following stocks: TH-Gal4^13^ (from Serge Birman, ESPCI, Université PSL, Paris); Ni-GFP, Dl-GFP, m3-GFP, Ser^Rx82^, UAS-Serrate (UAS-Ser) (from Schweisguth, Institut Pasteur, Paris); *MBSW* (Ron Davis, Scripps Institute, University of Florida), alrm-Gal4 (Marc Freeman, Vollum Institute, Oregon Health and Science University), 9-137-Gal4 (Ulrike Heberlein, Janielia farm Research Campus). Flies were either outcrossed at least two times to a reference strain (Cs), or to *w*^*-*^, to;*Sp*/*CyO*; to;;*Sb*/*TM6* previously outcrossed to Cs.

### Brain imaging

Unless otherwise stated 5-10 days old females flies were used. Flies were CO_2_-anesthetized and kept on ice until dissection.

#### GFP epifluorescence imaging and quantification in whole brains

Brains were quickly dissected in cold Phosphate-Buffered Saline (PBS) and fixed in 4% paraformaldehyde PBS for 10 minutes. They were then rinsed and mounted in PBS on polylysine-coated coverslips, and immediately imaged using a Leica epifluorescence microscope. Within an imaging session, the parameters were set on the most fluorescent brain and kept the same for all the conditions. The imageJ software (RRID:SCR_003070) was used to evaluate the background subtracted-average pixel intensity in the regions of interest. Except for the data presented in Figure 2, all the quantification were performed on the dorsal part of the brain, with a focus on the central complex, viewed from the posterior side. Experimenters were blind to the genotype when analyzing the images. The average pixel intensity in the control or reference condition was used to normalize the data and to combine several replicates of the same experiment. For each experiment, the control condition was dissected and analyzed together with flies belonging to one or several experimental conditions. Thus, the sample size of the control condition can be higher than the experimental conditions in several experiments.

#### Immunofluorescence in whole-mount brains

Brains were dissected in cold PBS, fixed for 20 min in a 4% paraformaldehyde PBS, washed in 0.3% Triton X-100 PBS (PBS-T) and blocked for 45 min in 5% goat serum in PBS-T, before overnight incubation in the corresponding primary antibody: rabbit anti-GFP (1:2000; Invitrogen, catalog #A6455; RRID:AB_221570), mouse anti-repo (DSHB Cat# 8D12 anti-Repo, RRID:AB_528448) and rabbit anti-TH (Millipore AB152). After primary antibody incubation, brains were washed in PBS-T and incubated with Alexa 488-conjugated and Alexa 633-conjugated secondary antibodies (Thermo Fisher Scientific, catalog A11008; RRID:AB_143165; catalog #A21050 RRID:AB_141431). Brains were mounted in Vectashield with DAPI (Vector Laboratories, catalog #H-1500; RRID:AB_2336788) and imaged using a confocal microscope (Zeiss LSM800). ImageJ was used for image processing. For repo-positive cell counting, identical imaging settings were applied across all genotypes. Optical section stacks were acquired at 1 μm intervals, spanning from the brain surface to the protocerebral bridge. Following background subtraction and Z-projection, cells were counted using the “Analyze Particles” function in ImageJ, with a minimum size threshold set at 2 μm in diameter. For TH immunofluorescence quantification, we quantified the average pixel intensity in the cell bodies of PPL1 neuons excluding the nuclear area.

### APS learning

5-10 day female flies were individually tested in the APS learning assay as previously described^14^. Briefly, each fly was placed in a T-maze and given a choice between entering a dark or a lighted vial. Naïve flies typically prefer the lighted vial, which in this assay is lined with filter paper soaked in quinine, creating an aversive association with light. Each fly underwent 16 trials over a period of 10–15 minutes. During the first 4 trials, flies generally exhibited photopositive behavior, but they progressively shifted toward photonegative choices over subsequent trials. The Learning Index (LI) was calculated as the difference in the number of photonegative choices made in the last 4 trials compared to the first 4 trials, thus a positive value indicates learning. This assay measures short-term memory, which lasts approximately 5 minutes, and is considered semi-operant^14^. Phototaxis was assessed using the same T-maze without quinine. 6 naïve flies were tested over 10 trials to evaluate their innate light preference. The photosensitive index was calculated as the percentage of photopositive choices. To assess quinine sensitivity, 6 naïve flies were individually inserted at the bottom of a uniformly lit 14 cm cylindrical tube. Each half of the tube was lined with filter paper—one soaked in quinine, the other dry. The quinine sensitivity index was calculated based on the amount of time a fly spent in seconds on the dry side over a 5-minute period. The experimenters were blind to the experimental conditions.

### Sleep recordings

3 days old females were individually inserted in 65mm long glass tubes filled with food at one end. Locomotor activity was recorded between age 5 days to 10 days using the DAMS infrared beam system from Trikinetics as previously described, with 12h:12h Light:Dark cycle. Sleep deprivation was carried out using the Sleep Nullifying Apparatus (SNAP) which elicits a negative geotaxis response every 10 seconds without eliciting stress response gene expression^15,16^.

### Drug feeding

For experiments involving the MBSW driver, RU486 (RU) (mifepristone, Sigma) was diluted in ethanol (50 mg/ml) and then diluted in melted standard food (100 mg/ml). Flies were fed RU486 for 5 days before dissection. For 3-Iodo-Tyrosine treatment, the drug was diluted 10mg/mL in melted standard food and fed to the flies for 6 days. L-DOPA was diluted 5mg/mL in standard melted food and fed to the flies for 24h, before dissection or learning evaluation, longer exposure led to learning impairments.

### Analysis of dopamine content using capillary UHPLC

5 brains per sample were quickly dissected from cold-anesthetized female flies and frozen in 5 µL of ringer solution at −80°C until being processed for tissue content analysis. 15 µL of ice-cold 0.1 mol/L perchloric acid containing 1.34 mmol/L EDTA and 0.05%, w/v sodium bisulfite were added to each sample and samples were then sonicated in an ultra-sonic bath for 12 min. The homogenates were centrifuged at 5,000 x g for 20 min at +4°C, and 10 µL of each supernatant was analyzed for monoamine content on the same day.

The UHPLC system consisted of a Prominence degasser, a LC-30 AD pump, a SIL-30AC autosampler, and a CTO-20AC column oven (Shimadzu). Detection was carried out at 40°C using a Decade II electrochemical detector fitted with a 0.7 mm glass carbon working electrode, an *in-situ* Ag/AgCl reference electrode, and a 25 m spacer (cell volume 80 nL, Antec, Leyden, The Netherlands). Separations were performed at 40°C (in detector oven) using a 100 × 0.32 mm Kappa Hypersil Gold 1.9 µm C18 column (Thermo Scientific). The mobile phase, which was pumped at 8.5 µL/min, consisted of 0.14 mol/L potassium phosphate, 0.1 mmol/L EDTA, 6 mmol/L OSA, 0.01% TEA (v/v), 8 mmol/L KCl, pH adjusted to 5 with 10 mmol/L sodium hydroxide, 6%methanol, and was filtered through a 0.22 m cellulose acetate membrane before use. The pressure was ~ 285 bars. Analytes were detected at an oxidation potential of 600 mV versus the reference electrode (range: 1nA, filter: 0.02 Hz). Chromatograms were acquired at a rate of 10 Hz using Lab Solutions software. The acquisition time was 30 min per sample and the retention time for DA was 14.1 min. On the day of analysis, the samples were placed in the autosampler and kept at +4°C before injection. The injection volume was 0.5 µL. Concentrations of dopamine in the extracts were determined by comparison of the chromatographic peak area with 10^-7^ mol/L of DA synthetic solution. Final concentrations of DA were expressed as pg per brain using the volume of extract (15 µL) and molar mass.

### Statistics

Distribution and homogeneity as well as statistical group comparisons were tested using Microsoft Excel plugin software Statel. Since the data were not normally distributed we used Man-Whitney (M-W) and Kruskall-Wallis (K-W) for statistical comparisons. In all cases averages with SEM are plotted. In the figures *: p<0,05, **: p<0,005, ***: p<0,0005, ****: p<0,00005.

## Results

We have previously reported that the Notch protein is expressed broadly in the brain, and that the canonical pathway is activated in a sub-population of glial cells, using the NRE-GFP reporter construct^17^. In this construct the DNA binding sequences of Su(H) are fused to a minimal promoter and to the GFP coding sequence. As seen on Figure 1 B-D, all the nuclei labelled by this construct are positive for the glial marker *reversed polarity* (*repo*). Labelled cells included a large subset of cortex-glial cells, as identified with the mesh like pattern of expression of GFP in the cortex. In addition, several surface, ensheathing and/or neuropil glial cells were also labelled. Accordingly, GFP could be detected around each neuropil and in a few arborizations within the calyx neuropil. We further evaluated the activation of canonical Notch signaling with the N-LV system, which is based on the expression of a chimeric receptor, where the intracellular domain is fused with LexA^18^. Using a LexAOP-GFP reporter construct combined with ubiquitous N-LexA expression, GFP immunofluorescence was detected in a subset of glial cells throughout the brain, a majority of which were cortex glial cells (Figure 1 E-G). Finally we also use the Ni-GFP line, where the wild type Notch receptor expression is replaced by a construct in which the intracellular domain is fused to GFP^19^ (Figure 1 H-J). As reported previously^19^, the nuclear translocation of the Notch intracellular domain is difficult to detect in *Drosophila*, but was nevertheless also observed in a subset of cortex-glial cells. Altogether, these results indicate that the Notch canonical pathway is activated in a subset of glial cells. In line with this conclusion, single cell transcriptomics data show a specific overlap of expression between *Repo, Notch*, and two targets of the Notch canonical pathway: *E(spl)mβ* and *E(spl)m3* (https://scope.aertslab.org)^20^. Indeed, flies expressing a CRISPR generated fusion of *m3* with GFP^21^ shows expression in a subset of glial cell nuclei (Figure 1 K-M). For the remaining of the study, we focused on the *NRE-GFP* reporter, which does not rely on the controlled expression of a chimeric receptor.

**Figure 1.**
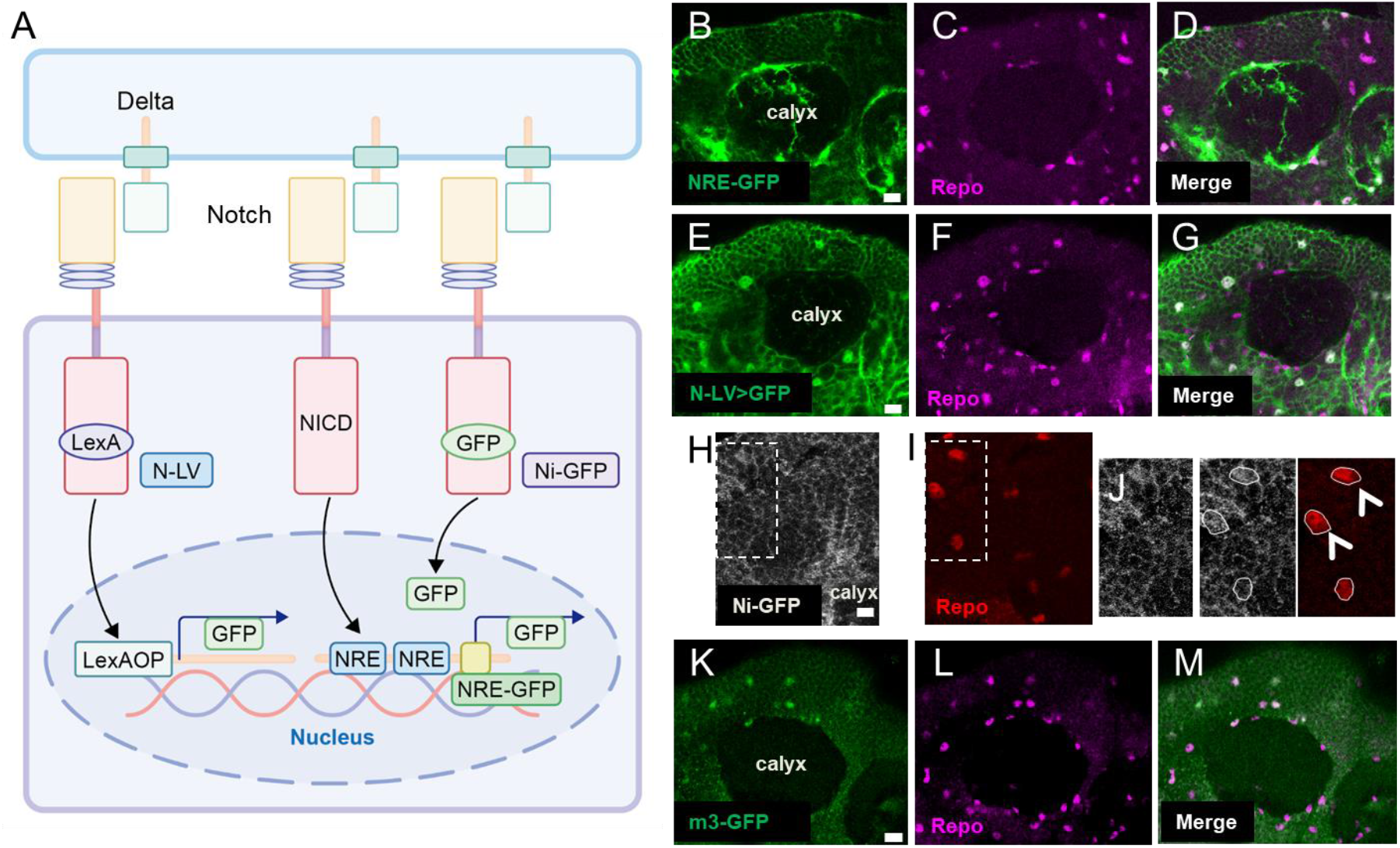
Diverse Notch canonical signaling reporters are expressed in a subset of glial cells. **A**, schematic of the different Notch reporter systems. **B-D**, *NRE-GFP* reporter; **E-G**, *N-LV>GFP* reporter; **H-J**, *Ni-GFP*, **I** shows enlargement of insets in **H** and **I**, colocalisation with repo in a subset of cortex glial cells (arrows). **K-M**, expression of CRISPR engineered *m3-GFP*. Single optical sections of the calyx showing immunofluorescent signal obtained with anti-GFP (green, left panels) and anti-repo (magenta or red, middle panels), scale bar: 10μm.

As detected with *NRE-GFP*, Notch activation was not uniform within the brain. Higher levels of expression were observed in the dorsal part of the brain, notably in the mushroom body lobes, the calices, the antennal lobes and the central complex (Figure 2). The ventral half of the central brain and the optic lobes showed the lowest levels of activation. In all those regions the labeled cells were glial.

**Figure 2.**
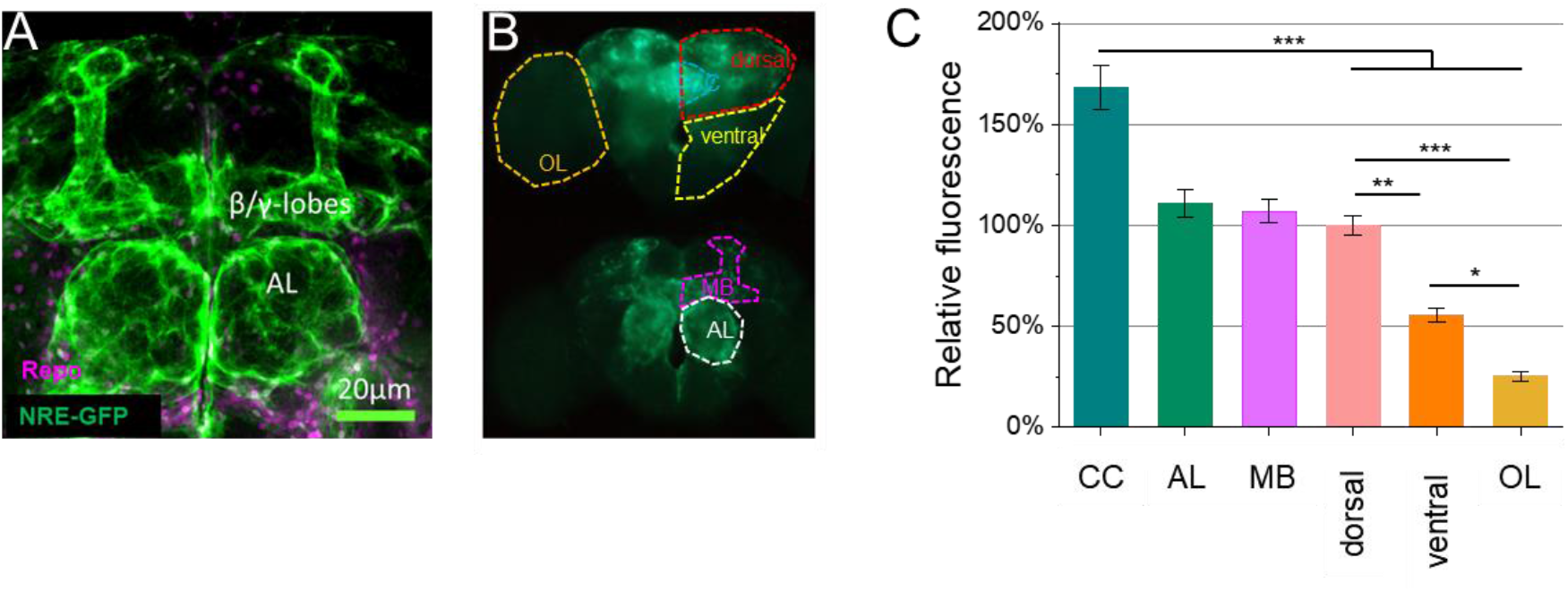
Notch canonical signaling is heterogeneously activated in the brain. **A**, confocal stack showing enhanced *NRE-GFP* expression in the mushroom body lobes and in the antennal lobes (AL). Anti-GFP immunofluorescence is shown in green, anti-repo immunofluorescence in magenta. **B**, whole brain images obtained with an epifluorescent microscope, showing endogenous GFP fluorescence in the posterior (top) and frontal (bottom) part of the brain. OL: optic lobe, AL: antennal lobe, CC central complex. **C**, quantification of the GFP fluorescence in the different brain areas, normalized to the dorsal part. K-W, p<0,00001, n=12-25.

To further confirm that the *NRE-GFP* expression was dependent on Notch, we used the Gal4/UAS system to drive Notch transgenes to down- or up-regulate the pathway. Expressing Notch transgenes with the pan glial repo-Gal4 driver was lethal, probably due to developmental defects. Using the Eaat1-Gal4 driver, and several more specific drivers (see below) we were able to obtain viable adults. The Eaat1-Gal4 driver contains the regulatory sequences of the glutamate transporter Eaat1^22^ and is expressed in astrocyte-like, ensheathing and cortex glia^23^. Knockdown of Notch using *Eaat1-Gal4* to drive a UAS-RNAi transgene resulted in a partial to near-complete loss of *NRE-GFP* expression (Figure 3 A-B). Conversely, expressing the intracellular domain of Notch in the same cells resulted in an almost 150% upregulation of the reporter. Notch signaling is involved in gliogenesis, and those changes in *NRE-GFP* expression may originate from a change in the number of glial cells. However, immunostaining for Repo revealed no change in the number of glial cell nuclei in both gain and loss of function conditions (Figure 3 C), indicating that the changes in *NRE-GFP* expression was not due to developmental defects or glial cell loss. Furthermore, temporally restricting *Notch* knockdown to the adult stage using the TARGET system^24^ also resulted in a significant, though less pronounced, reduction in *NRE-GFP* expression (Figure 3 D). Altogether these results suggest that the majority of canonical Notch signaling occurs in Eaat1-expressing glial cells. Supporting this conclusion, downregulation of *Su(H)* in the same glial population led to an approximate 50% reduction in *NRE-GFP* expression (Figure 3 E). To determine which of the known Notch ligands, Delta (Dl) or Serrate (Ser), was involved in Notch activation in this context we evaluated NRE-GFP expression in flies heterozygous for *Dl* or *Ser* loss of function alleles (Figure 4 A). *Dl* heterozygous mutant flies showed a significant reduction of reporter expression, whereas no detectable downregulation was observed in *Ser* heterozygous mutant flies, suggesting that Dl is the main Notch ligand. In line with this result, single cell transcriptomics data (https://scope.aertslab.org)^20^ show *Dl* expression but no detectable expression for *Ser* in the brain except in a very small group of cells that appear to be hemocytes (based on co-expression of marker genes such as Serpent, Nimrod, CG8501). None of the UAS-Dl-RNAi transgenes that we tested with the pan-neuronal *elav-Gal4* or mushroom body specific *MBSW* driver provided a reduction in *NRE-GFP* expression (see methods), potentially because of the lack of knockdown efficiency. In contrast, we found that overexpression of *Dl* in the mushroom bodies resulted in a 300% increase in *NRE-GFP* expression in the dorsal part of the brain, around the calices and the lobes, whereas overexpression of *Ser* had no effect (Figure 4 B-C). Interestingly, driving *Dl* expression in glial cells using the Eaat1-Gal4 driver also resulted in a strong induction of the *NRE-GFP* reporter, throughout the brain, including the optic lobes, where NRE-GFP is normally almost undetectable (Figure 4 D-E). We previously reported Dl expression in neurons by immunostaining. To further characterize Dl expression in the brain we used a CRISPR-generated GFP tagged allele of the gene^25^. As shown in Figure 4F, we could confirm Dl expression in the adult brain using this tool. Dl was primarily detected in the cell bodies of the cortex, with significantly lower expression observed in the neuropil regions, in agreement with previous data.

**Figure 3.**
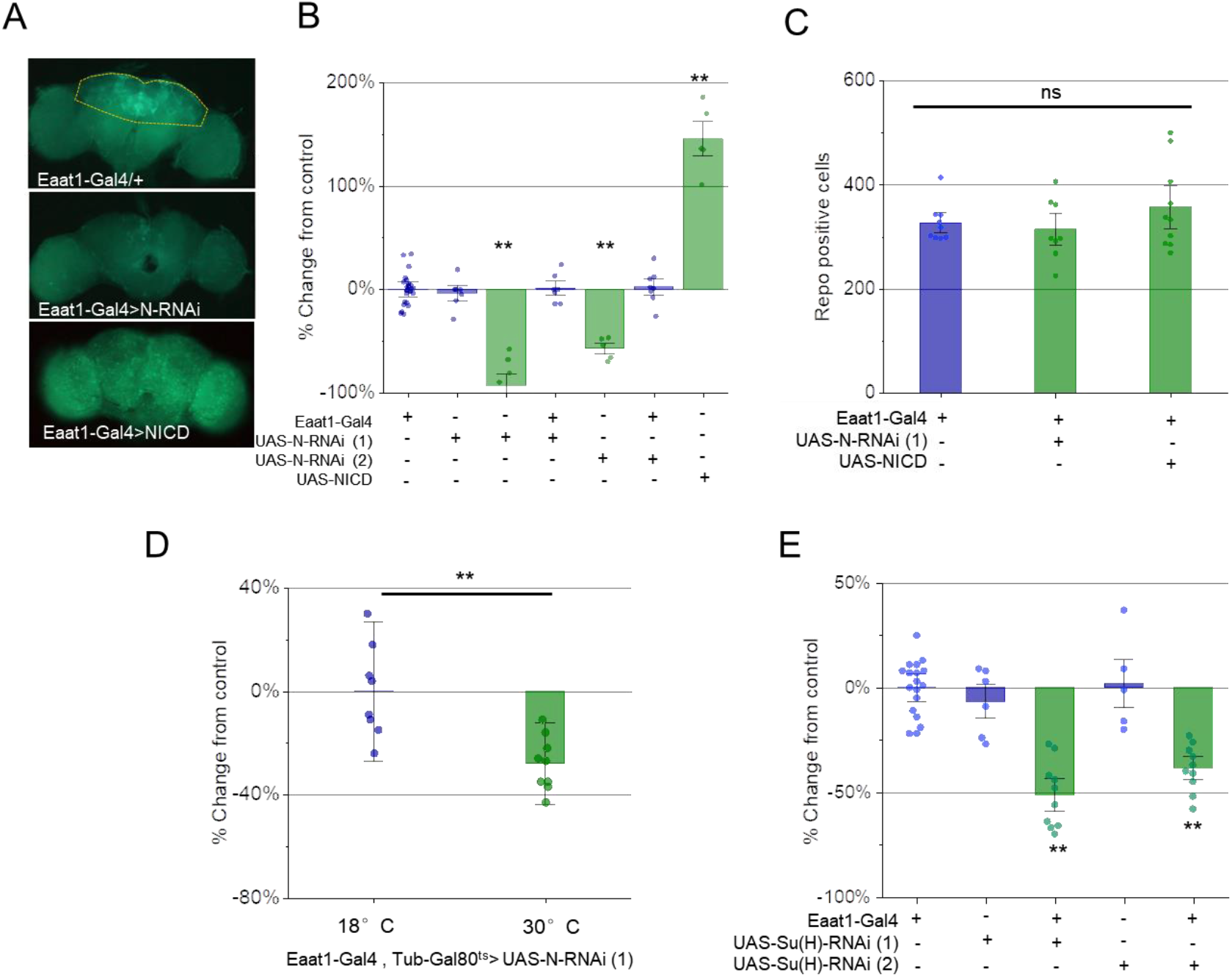
Notch canonical signaling at the adult stage relies on Su(H). **A**, whole brain images obtained with an epifluorescent microscope, showing endogenous GFP fluorescence in the posterior part of the brain. The doted line in the top image shows the area quantified in B. Expression of an UAS-N-RNAi transgene (in this case *UAS-N-RNAi (1)*) noticeably reduces *NRE-GFP* expression, while expression of the NICD results in induction of the reporter throughout the brain. **B**, quantification of the GFP fluorescence upon Notch knockdown in glia or expression of the NICD using the *Eaat1-Gal4* driver. K-W: p<0,00001, n=5-7 (experimental) and 27 (*Eaat1-Gal4* controls). * refers to comparison with Gal4 and UAS alone control groups. **C**, quantification of the number of repo-positive cells in the dorsal area of the brain using anti-repo immunofluorescence (see methods), in the conditions shown in A-B. K-W, p<0,00001, n=5-7 (experimental) and 27 (*Eaat1-Gal4* controls). K-W: p<0,51, n=9-10 brains. **D**, quantification of the GFP fluorescence upon knockdown of *Notch* at the adult stage in glia using the TARGET system. Flies were transferred to 30°C for 10 days and compared to flies maintained at 18°C. Flies were raised at 18°C. M-W: p<0,0001, n=8-9. **E**, quantification of the GFP fluorescence upon knockdown of *Su(H)*. K-W: p<0,00001, n=5-10 (experimental) and 18 (*Eaat1-Gal4* controls). * refers to comparison with Gal4 and UAS alone control groups.

**Figure 4.**
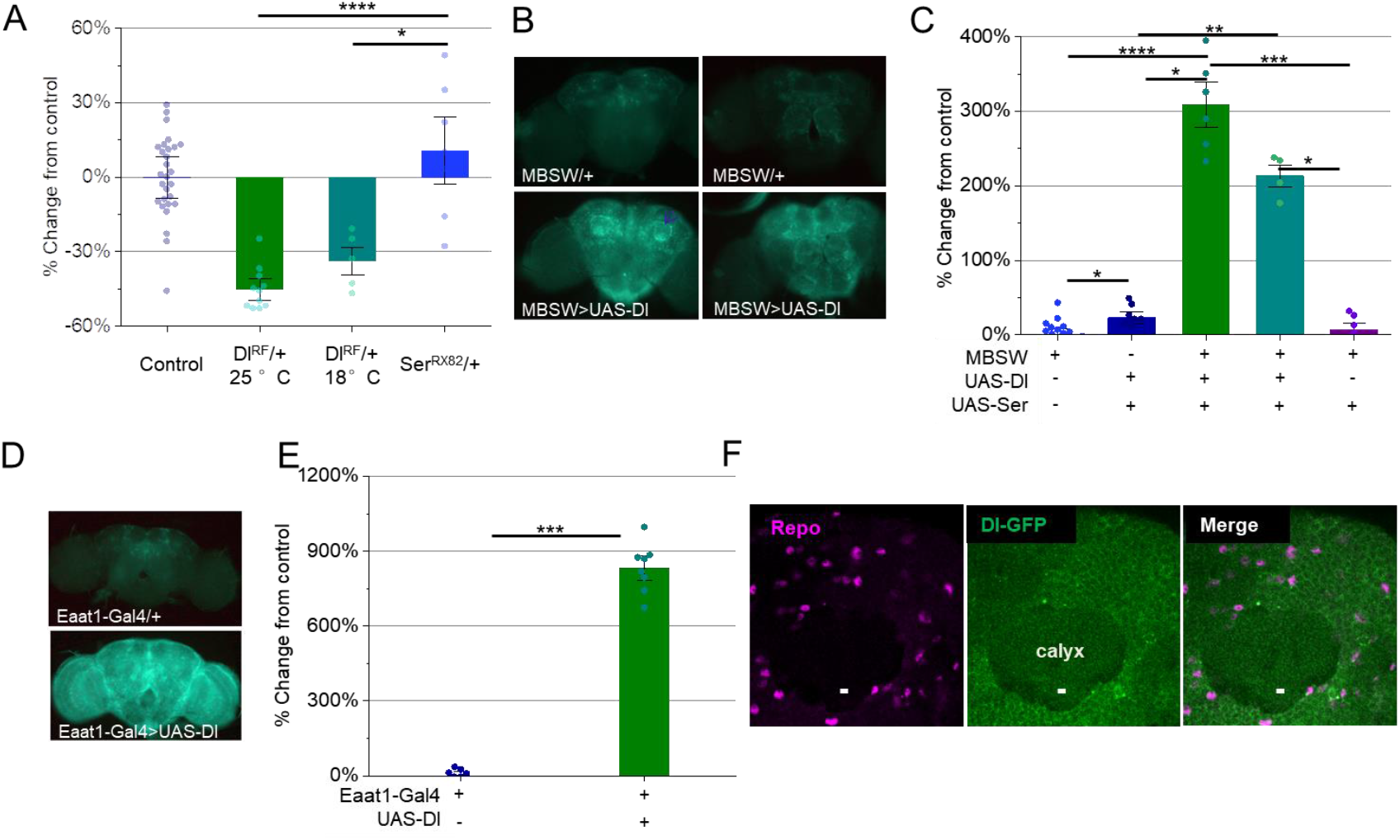
Notch canonical signaling in glia relies on the Delta ligand. **A**, quantification of the GFP fluorescence in *NRE-GFP* bearing flies heterozygous for the *Dl*^*RF*^ *or Ser*^*RX82*^mutants. Since *Dl*^*RF*^ has been reported to be thermosensitive, a subset of flies were raised and maintained at 18°C. K-W: p<0,00001, n=5-11 (experimental) and 28 (Eaat1-Gal4 controls). **B**, whole brain images obtained with an epifluorescent microscope, from the posterior (left) and frontal (right) sides, showing increased expression of *NRE-GFP* notably in the calyx (arrow) and the mushroom body lobes. **C**, quantification of the GFP fluorescence upon overexpression of *Dl* or *Ser* in the mushroom body neurons using the RU-inducible *MBSW* driver. All flies were fed RU except the “no RU” group,revealing leaky expression of the driver. K-W: p<0,00001, n=4-8 (experimental) and 22 (*MBSW* controls). **D**, whole brain images obtained with an epifluorescent microscope, from the posterior side, showing strong enhancement of *NRE-GFP* expression throughout the brain, upon Dl induced expression in glia using the *Eaat1-Gal4* driver. **E**, quantification of the GFP fluorescence upon driving *Dl* expression in *Eaat1-Gal4*-positive cells. M-W: p<0,0003 n=7-8. **F**, Expression of a CRISPR engineered *Dl-GFP*, single optical sections of the calyx with immunofluorescent signal obtained with anti-GFP (green, left panels) and anti-repo (magenta or red, middle panels), scale bar: 5μm.

We then used the *NRE-GFP* reporter to assess how Notch signaling was influenced by various conditions affecting brain plasticity and learning. As shown in Figure 5 A-B we observed a progressive downregulation of reporter expression during the first four days of adult life, followed by stabilization once the flies reached maturity (5 days). Interestingly, *NRE-GFP* is upregulated following social enrichment in young male flies (Figure 5 C-D), suggesting that it is associated with higher sleep need and/or brain plastic processes^26^ involved in this assay. In contrast, a 12h sleep deprivation during the night led to a reduction of *NRE-GFP* expression (Figure 5 E).

**Figure 5.**
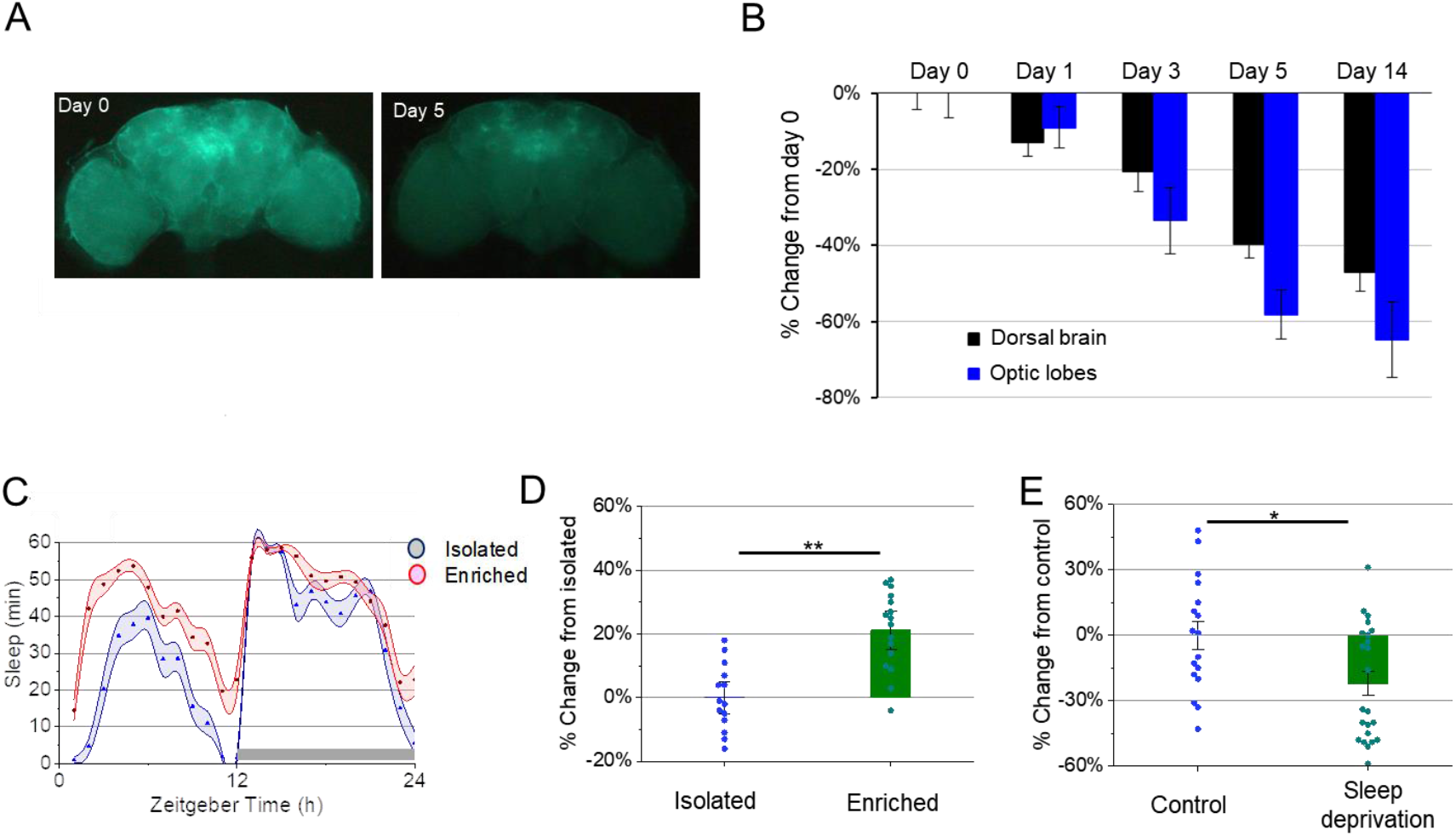
Notch canonical signaling is modulated by age, neuronal plasticity and sleep deprivation. **A**, whole brain images obtained with an epifluorescent microscope, from the posterior side showing *NRE-GFP* expression at day 0 and day 5. **B**, quantification of the GFP fluorescence in the dorsal brain and in the optic lobes at different ages. K-W: p<0,00001, n=5-12 (day1-14) and 30 (day0). **C**, 24h sleep curve, for *NRE-GFP* expressing male flies either isolated at the adult stage (“isolated”, open diamonds) or maintained in a group of 64 individuals during 5 days (“enriched”, black squares). The black bar indicates the dark phase. Higher sleep amounts are observed in the enriched condition as previously reported. **D**, quantification of the GFP fluorescence in the enriched condition compared to isolated. M-W: p<0,00025, n=14-16. **E**, quantification of the GFP fluorescence in sleep deprived (12h, from ZT12 to ZT0) *NRE-GFP* expressing flies.

We previously demonstrated that constitutive activation of Notch signaling in glia—via expression of the Notch intracellular domain—reduces the vulnerability of APS to sleep deprivation^7^. To further investigate the role of Notch signaling in APS learning, we assessed flies with Notch knockdown. As shown in Figure 6 A-B, downregulation of *Notch* impaired APS learning, while leaving phototaxis and quinine sensitivity intact (Figure 6 D). A similar phenotype was observed with knockdown of *Su(H)*, indicating that the canonical Notch pathway is required for this form of learning (Figure 6 D,C). None of these two knockdown conditions significantly impacted nighttime sleep quota, or average nighttime sleep bout duration (Figure 6 E-F), which are known to affect APS performance^2^, suggesting that Notch signaling directly affects the learning process. We next targeted specific glial subtypes—including surface glia, cortex glia, ensheathing glia, and astrocyte-like glia—to identify which populations are critical for APS learning (Figure 6 G-H). Knockdown of *Notch* in surface-cortex- or ensheathing glia was sufficient to impair APS learning, while sparing phototaxis and quinine sensitivity (Figure 6 H-I). In contrast, knockdown in astrocyte-like glia had no detectable effect, in line with the low expression of the NRE-GFP expression in those cells.

**Figure 6.**
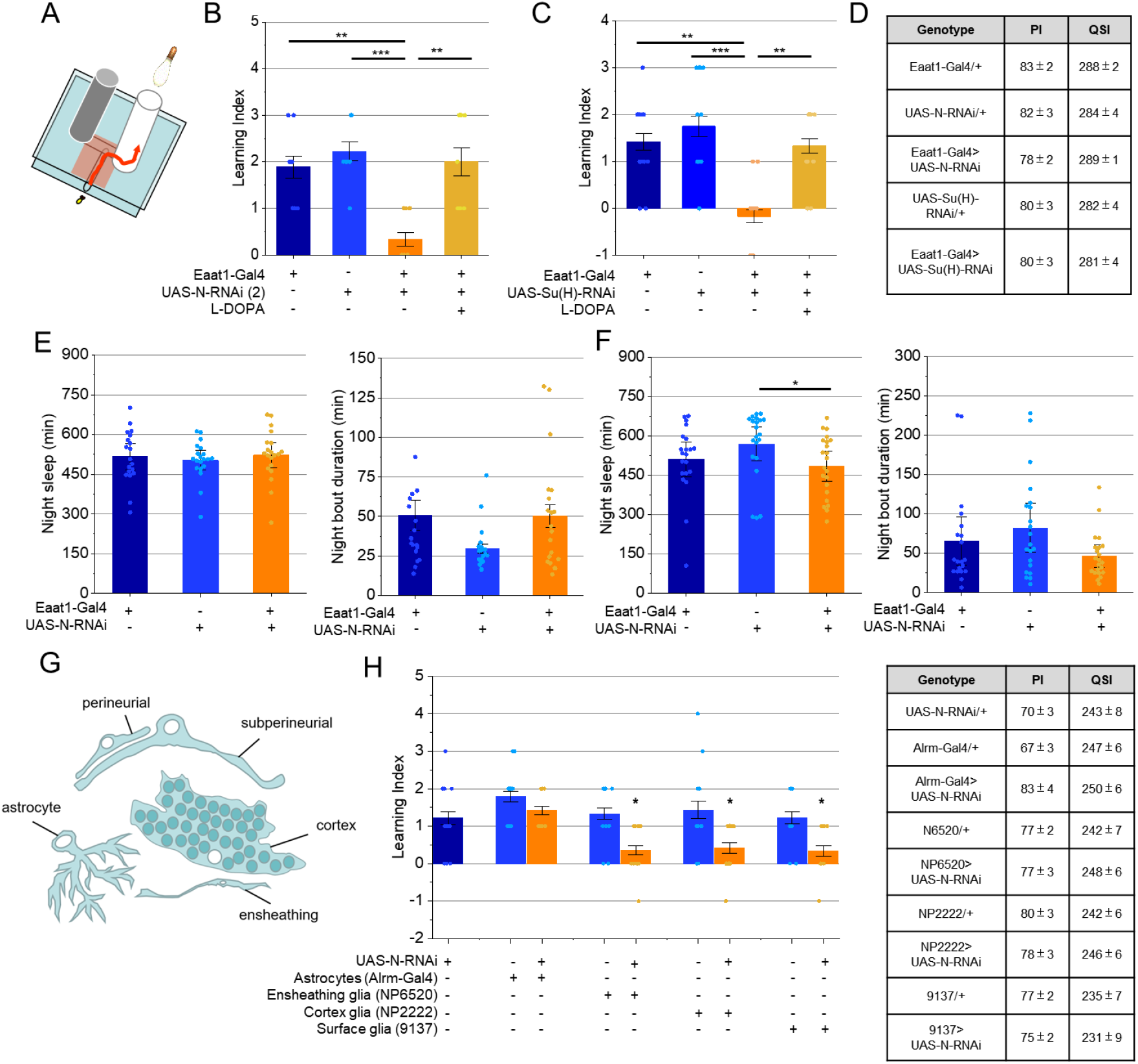
Notch canonical signaling in glia is required for APS learning. **A**, schematic of the learning assay. **B-C and G**, evaluation of short-term memory in the APS assay, a high Learning Index indicates successful learning (formation of short-term memory). **B**, knockdown of *Notch* in *Eaat1-Gal4* positive cells impairs learning. Feeding flies L-DOPA rescues normal learning. K-W P<0,0005, n=9 for all groups. **C**, knockdown of *Su(H)* in *Eaat1-Gal4* positive cells impairs learning. Feeding flies L-DOPA rescues normal learning performance. K-W P<0,0002, n=12 for all groups. **D**, sensory metrics: Phototaxis Index (PI) and Quinine Sensitivity Index (QSI) are similar in all the genotypes (n=6). **E-F**, total night sleep (left) and average night bout sleep duration (right) are similar in knockdown and control flies. E, K-W: p< (left), K-W: p<0,07 (right), n=19-20. F, K-W: p<0,023 (left), K-W: p<0,19 (right), n=20-22. **G**, schematic of the different glial cell types in the *Drosophila* brain. **H**, evaluation of short-term memory using the APS assay upon knockdown of *Notch* in different glial cell types. K-W: p<0,00002, n=9-14. * refers to comparison with Gal4 and UAS alone control groups. **I**, sensory metrics are similar in all the genotypes (n=6).

APS relies on dopaminergic transmission^2^ and remarkably, the learning deficits in both *Notch* and *Su(H)* knockdown flies were fully rescued by feeding L-DOPA (Figure 6 B-C) showing that enhancing dopaminergic transmission can compensate for impaired Notch signaling. These results led us to investigate the relationship between Notch signaling and dopaminergic transmission. We first assessed the impact of 3-iodo-tyrosine or L-DOPA feeding to respectively down- or up-regulate brain dopamine levels (Figure 7 A).Dopamine downregulation led to a moderate reduction of *NRE-GFP* expression, similar to the one observed after sleep deprivation. In contrast, L-DOPA feeding did not significantly affect the expression of the reporter. We also measured whole-brain dopamine levels by HPLC and found no significant changes in either Notch knockdown flies or those expressing the Notch intracellular domain (Figure 7 B-C). In line with these findings, tyrosine hydroxylase levels assessed by immunofluorescence in PPL1 neurons, that mediate aversive that mediate aversive reinforcement^27^, were unaffected by downregulation of Notch in Glia (Normalized pixel intensity, Eaat1-Gal4/+: 100±2%, Eaat1-Gal4>UAS-N-RNAi (1): 102±2%, 11-15 brains and 175-202 neurons). These results suggest that Notch signaling does not significantly influence dopaminergic transmission, whereas dopaminergic transmission can modulate Notch signaling. This is consistent with prior observations suggesting that sleep deprivation downregulates both Notch signaling (^7^, Figure 5E) and dopaminergic activity in APS-sensitive circuits^2^. To further test this idea, we overexpressed Dl in dopaminergic neurons during sleep deprivation, in order to activate Notch in the direct vicinity of those neurons. As expected, sleep deprivation impaired learning in control flies (Figure 7 D-E). However, *Dl* overexpression preserved normal learning performance without affecting phototaxis or quinine sensitivity. In line with this result, we detect Dl expression in PPL1 dopaminergic neurons (Figure 7F).

**Figure 7.**
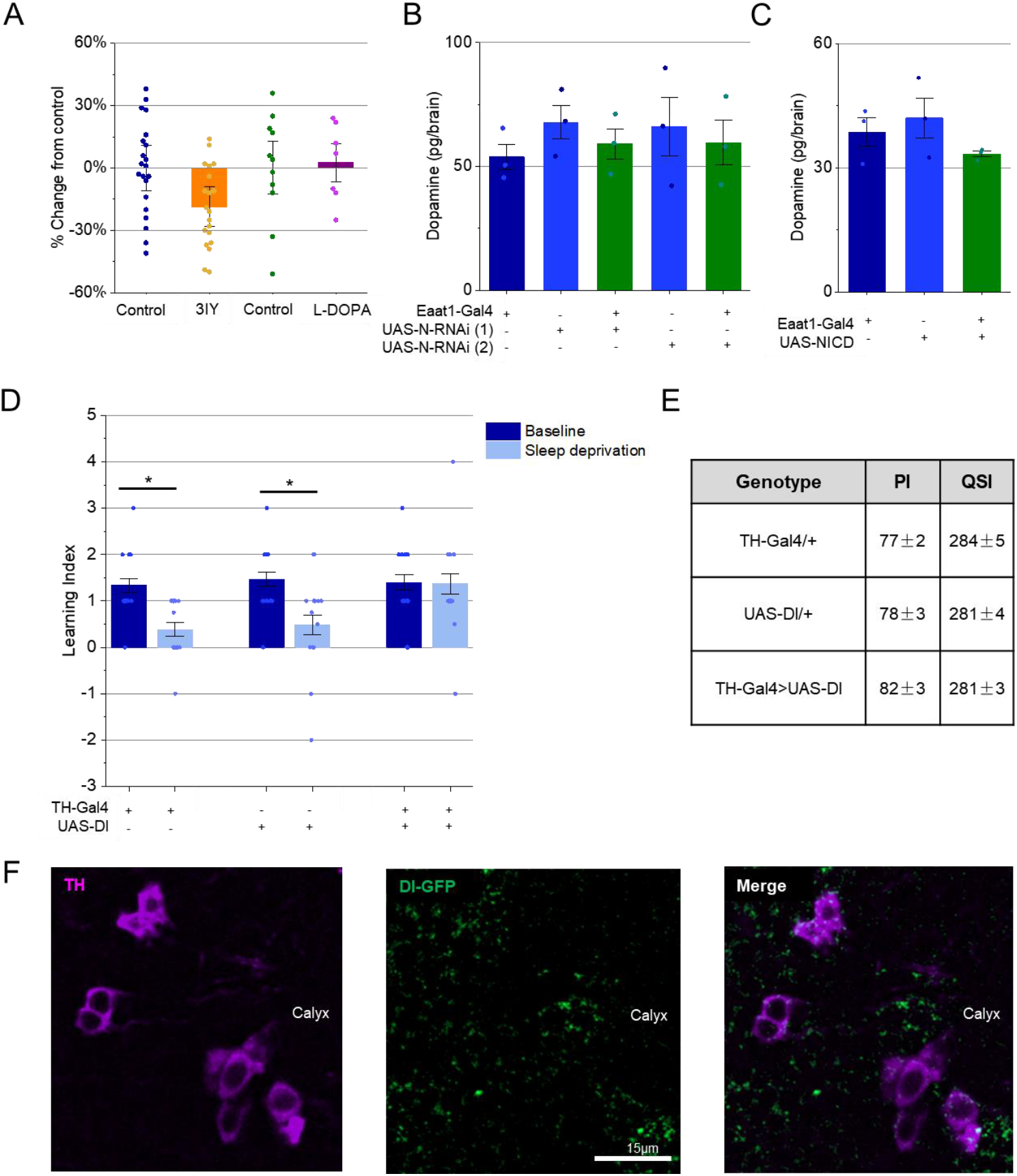
Notch canonical signaling in glia is modulated by dopaminergic transmission. **A**, quantification of the GFP fluorescence in *NRE-GFP* expressing flies, fed 3IY or L-DOPA, compared to their respective controls. M-W (3IY): p<0,011, n=20-21. M-W (L-DOPA): p<1, n=7-11. **B-C**, HPLC quantification of dopamine in the brains of flies knockdown for *Notch* (**B**) or expressing NICD (**C**) using the *Eaat1-Gal4* driver (n=3). **D**, evaluation of short-term memory using the APS assay upon in flies overexpressing *Dl* in dopaminergic neurons using the TH-Gal4 driver. Control flies show impaired performance after a 12h sleep deprivation, while Dl overexpressing flies maintain normal learning. M-W (*TH-Gal4*): p<0,0035, n=15, M-W (*UAS-Dl*): p<0,036, n=12-15, M-W (*TH-Gal4>UAS-Dl*): p<0,82, n=15. **E**, sensory metrics are similar in all the genotypes tested (n=6). Sleep deprivation does not affect sensory metrics^2^. **F**, PPL1 dopaminergic neurons, revealed by anti-THimmunoreactivity (magenta, left), and expressing Dl-GFP (green, center).

## Discussion

Here, we show that canonical Notch signaling is activated in a specific subset of glial cells, primarily cortex and ensheathing glia. This signaling is essential for the formation of short-term memory in the aversive phototaxic suppression (APS) assay, which is sensitive to both sleep deprivation and to dopaminergic transmission. We find that reduced dopaminergic transmission leads to a downregulation of Notch signaling in these glial cells. Given that sleep deprivation is known to impair dopaminergic transmission in memory-relevant neuronal circuits, our results suggest a potential mechanism by which sleep deprivation leads to learning deficits—through the disruption of dopamine-dependent Notch signaling in glia.

While there are very few reports on the role of Notch signaling in sleep regulation and short-term memory, numerous studies in both *Drosophila* and murine models have investigated its function in the adult brain, particularly in relation to memory and other forms of long-term neuronal plasticity^28,29^. Most of this research has focused on Notch activity within neurons or employed genetic tools that affect the pathway ubiquitously^3,4,9–11,30^. In *Drosophila*, both canonical and non-canonical Notch signaling pathways have been implicated in long-term plasticity, particularly within the mushroom bodies and the olfactory system^5,10,29^. In contrast, our findings suggest that canonical Notch signaling is primarily active in glial cells. This observation is consistent with single-cell transcriptomic data from both Drosophila^20^ and mouse brains^12^. Thus, Notch canonical signaling may be activated relatively parsimoniously in neurons compared to glia, and play different roles in each cell type.

What could be the function of Notch canonical signaling in this particular context? This study suggests that ensheathing, cortex and surface glia are all involved. Ensheathing glia enwraps each neuropil, providing a shielded environment for axons, dendrites and synapses^31,32^ and a diffusion barrier that may facilitate neurotransmission^33^. Cortex glial cells envelop all the neuronal cell bodies, which are preferentially located in the most superficial layers of the brain in *Drosophila*. A single cortex glial cell can surround up to 100 neuronal cell bodies and their associated neurites as they extend towards the neuropil, thus constituting a trophospongium^31^. They are also closely associated with the subperineurial glia, which together with the more peripheral perineurial cells constitute the *Drosophila* equivalent of the blood brain barrier, providing nutrients to the brain^34^, and protecting it from the hemolymph. Given these morphological features, and the low levels of Delta expression in the neuropil (^7^ and this report) it appears relatively unlikely that Notch is directly modulating neuronal plasticity at the level of the relevant synapses. Based on the reporters examined in this study, Notch canonical signaling is upregulated in the mushroom bodies, the central complex and in olfactory glomeruli, structures that are known to display neuronal plasticity, and at least for the former two, to regulate sleep. Plastic processes in those learning associated structures may upregulate Dl expression in neurons, leading to enhanced Notch activation in the surrounding glial cells and to the subsequent expression of transcriptional targets that may be permissive for the formation of short-term memory. Ensheathing glial cells have been shown to affect behavior via a metabolic role^35^ or as a relay of aversive sensory input^33^. The latter function may be relevant for APS, since ensheathing glia rely on dopaminergic signaling in this context^33^. While ensheathing glia can envelop entire neuropiles, the specific effects on synapses appear to be driven by neuronal cues^33^. The involvement of a similar regulatory mechanism in Notch signaling could account for the pronounced activation of the pathway in ensheathing glia adjacent to neuropils associated with high plasticity (Figure 2). Cortex glia regulate neuronal activity at the neuromuscular junction through Ca^2+^ signaling^36^. If such a function is maintained in the adult fly, it may prove crucial for the formation of short-term memory, and would be a prime candidate for future investigations regarding *Notch* function. Interestingly, a recent report has shown that surface glia, in particular perineurial glia, could also regulate neuronal activity through Ca^2+^ waves^37^. The regulation of neuronal activity by glial Ca^2+^ could implicate neuronal metabolism. In the larva, synchronized Ca^2+^ waves in subperineurial glia have been shown to regulate nutrient import^38^. Besides, cortex glia can provide key energy metabolites linked to the formation of long-term memory^39,40^, and a similar type of mechanism may be required for learning and the regulation of sleep homeostasis. Cortex glia and ensheathing glia are involved in neurotransmitter recycling ^23,41–43^, which may also be of particular importance for learning and for sleep homeostasis^43^. Recent studies have shown that astrocytic calcium signaling regulates neuronal activity and sleep homeostasis^44–46^. Although current findings suggest that Notch signaling primarily operates through non-astrocytic glial cell types, interactions among various glial populations likely contribute to neuronal activity, synaptic plasticity, and sleep-wake cycle regulation^47^.

We report here that sleep deprivation or inhibiting dopamine synthesis by feeding flies 3-iodo-tyrosine leads to a downregulation of Notch signaling in glia. We have previously shown that sleep deprivation impairs dopaminergic transmission in the mushroom bodies, which is required for the APS learning, and reduced *Dl* mRNA expression^7^. These results thus suggest that learning impairments associated with sleep deprivation may originate, at least in part, from a downregulation of Notch signaling in response to disrupted dopaminergic transmission. In agreement with this idea, upregulation of Notch signaling near dopaminergic neurons can prevent learning impairments associated with sleep deprivation. Dopamine levels were unaffected by Notch signaling in glia, indicating that Notch is acting downstream of dopaminergic transmission. Interestingly, L-DOPA supplementation restored learning in Notch and Su(H) knockdown flies, suggesting that dopaminergic signaling can compensate for impaired learning through a Notch-independent mechanism.

## Acknowledgments

This work was funded by INSERM (https://www.inserm.fr), CNRS (https://www.cnrs.fr/fr/page-daccueil), Université Claude Bernard Lyon 1 (https://www.univ-lyon1.fr), région Auvergne-Rhône-Alpes and SFRMS (www.sfrms-sommeil.org). HL received funding from National Natural Science Foundation of China (Grant # 82401744).

## References

(1) Heitzler, P. Biodiversity and Noncanonical Notch Signaling. Current topics in developmental biology 2010, 92, 457–481. 10.1016/S0070-2153(10)92014-0.

(2) Seugnet, L.; Suzuki, Y.; Vine, L.; Gottschalk, L.; Shaw, P. J. D1 Receptor Activation in the Mushroom Bodies Rescues Sleep-Loss-Induced Learning Impairments in Drosophila. Curr Biol 2008, 18 (15), 1110–1117. 10.1016/j.cub.2008.07.028.

(3) Ge, X.; Hannan, F.; Xie, Z.; Feng, C.; Tully, T.; Zhou, H.; Zhong, Y. Notch Signaling in Drosophila Long-Term Memory Formation. Proc Natl Acad Sci U S A 2004, 101 (27), 10172–10176.

(4) Presente, A.; Boyles, R. S.; Serway, C. N.; de Belle, J. S.; Andres, A. J. Notch Is Required for Long-Term Memory in Drosophila. Proc Natl Acad Sci U S A 2004, 101 (6), 1764–1768.

(5) Kidd, S.; Lieber, T. Mechanism of Notch Pathway Activation and Its Role in the Regulation of Olfactory Plasticity in Drosophila Melanogaster. PloS one 2016, 11 (3), e0151279. 10.1371/journal.pone.0151279.

(6) Kidd, S.; Struhl, G.; Lieber, T. Notch Is Required in Adult Drosophila Sensory Neurons for Morphological and Functional Plasticity of the Olfactory Circuit. PLoS Genet 2015, 11 (5), e1005244. 10.1371/journal.pgen.1005244.

(7) Seugnet, L.; Suzuki, Y.; Merlin, G.; Gottschalk, L.; Duntley, S. P.; Shaw, P. J. Notch Signaling Modulates Sleep Homeostasis and Learning after Sleep Deprivation in Drosophila. Curr Biol 2011, 21 (10), 835–840. 10.1016/j.cub.2011.04.001.

(8) Singh, K.; Chao, M. Y.; Somers, G. A.; Komatsu, H.; Corkins, M. E.; Larkins-Ford, J.; Tucey, T.; Dionne, H. M.; Walsh, M. B.; Beaumont, E. K.; Hart, D. P.; Lockery, S. R.; Hart, A. C. C. Elegans Notch Signaling Regulates Adult Chemosensory Response and Larval Molting Quiescence. Current biology : CB 2011, 21 (10), 825–834. 10.1016/j.cub.2011.04.010.

(9) Zhang, J.; Yin, J. C.; Wesley, C. S. Notch Intracellular Domain (NICD) Suppresses Long-Term Memory Formation in Adult Drosophila Flies. Cellular and molecular neurobiology 2015, 35 (6), 763–768. 10.1007/s10571-015-0183-9.

(10) Zhang, J.; Little, C. J.; Tremmel, D. M.; Yin, J. C.; Wesley, C. S. Notch-Inducible Hyperphosphorylated CREB and Its Ultradian Oscillation in Long-Term Memory Formation. The Journal of neuroscience : the official journal of the Society for Neuroscience 2013, 33 (31), 12825–12834. 10.1523/JNEUROSCI.0783-13.2013.

(11) Gerstner, J. R.; Vanderheyden, W. M.; Shaw, P. J.; Landry, C. F.; Yin, J. C. P. Fatty-Acid Binding Proteins Modulate Sleep and Enhance Long-Term Memory Consolidation in Drosophila. PLOS ONE 2011, 6 (1), e15890. 10.1371/journal.pone.0015890.

(12) Cahoy, J. D.; Emery, B.; Kaushal, A.; Foo, L. C.; Zamanian, J. L.; Christopherson, K. S.; Xing, Y.; Lubischer, J. L.; Krieg, P. A.; Krupenko, S. A.; Thompson, W. J.; Barres, B. A. A Transcriptome Database for Astrocytes, Neurons, and Oligodendrocytes: A New Resource for Understanding Brain Development and Function. J Neurosci 2008, 28 (1), 264–278. 10.1523/JNEUROSCI.4178-07.2008.

(13) Riemensperger, T.; Isabel, G.; Coulom, H.; Neuser, K.; Seugnet, L.; Kume, K.; Iché-Torres, M.; Cassar, M.; Strauss, R.; Preat, T.; Hirsh, J.; Birman, S. Behavioral Consequences of Dopamine Deficiency in the Drosophila Central Nervous System. Proc Natl Acad Sci U S A 2011, 108 (2), 834–839. 10.1073/pnas.1010930108.

(14) Seugnet, L.; Suzuki, Y.; Stidd, R.; Shaw, P. J. Aversive Phototaxic Suppression: Evaluation of a Short-Term Memory Assay in Drosophila Melanogaster. Genes Brain Behav 2009, 8 (4), 377–389. 10.1111/j.1601-183X.2009.00483.x.

(15) Shaw, P. J.; Cirelli, C.; Greenspan, R. J.; Tononi, G. Correlates of Sleep and Waking in Drosophila Melanogaster. Science 2000, 287 (5459), 1834–1837. 10.1126/science.287.5459.1834.

(16) Shaw, P. J.; Tononi, G.; Greenspan, R. J.; Robinson, D. F. Stress Response Genes Protect against Lethal Effects of Sleep Deprivation in Drosophila. Nature 2002, 417 (6886), 287–291. 10.1038/417287a.

(17) Stempfle, D.; Kanwar, R.; Loewer, A.; Fortini, M. E.; Merdes, G. In Vivo Reconstitution of Gamma-Secretase in Drosophila Results in Substrate Specificity. Mol Cell Biol 2010, 30 (13), 3165–3175. 10.1128/MCB.00030-10.

(18) Lieber, T.; Kidd, S.; Struhl, G. DSL-Notch Signaling in the Drosophila Brain in Response to Olfactory Stimulation. Neuron 2011, 69 (3), 468–481. 10.1016/j.neuron.2010.12.015.

(19) Couturier, L.; Vodovar, N.; Schweisguth, F. Endocytosis by Numb Breaks Notch Symmetry at Cytokinesis. Nat Cell Biol 2012, 14 (2), 131–139. 10.1038/ncb2419.

(20) Davie, K.; Janssens, J.; Koldere, D.; De Waegeneer, M.; Pech, U.; Kreft, Ł.; Aibar, S.; Makhzami, S.; Christiaens, V.; Bravo González-Blas, C.; Poovathingal, S.; Hulselmans, G.; Spanier, K. I.; Moerman, T.; Vanspauwen, B.; Geurs, S.; Voet, T.; Lammertyn, J.; Thienpont, B.; Liu, S.; Konstantinides, N.; Fiers, M.; Verstreken, P.; Aerts, S. A Single-Cell Transcriptome Atlas of the Aging Drosophila Brain. Cell 2018, 174 (4), 982–998.e20. 10.1016/j.cell.2018.05.057.

(21) Lleo, A.; Berezovska, O.; Ramdya, P.; Fukumoto, H.; Raju, S.; Shah, T.; Hyman, B. T. Notch1 Competes with the Amyloid Precursor Protein for Gamma-Secretase and down-Regulates Presenilin-1 Gene Expression. J Biol Chem 2003, 278 (48), 47370–47375. 10.1074/jbc.M308480200.

(22) Rival, T.; Soustelle, L.; Strambi, C.; Besson, M. T.; Iche, M.; Birman, S. Decreasing Glutamate Buffering Capacity Triggers Oxidative Stress and Neuropil Degeneration in the Drosophila Brain. Curr Biol 2004, 14 (7), 599–605.

(23) Farca Luna, A. J.; Perier, M.; Seugnet, L. Amyloid Precursor Protein in Drosophila Glia Regulates Sleep and Genes Involved in Glutamate Recycling. J Neurosci 2017, 37 (16), 4289–4300. 10.1523/JNEUROSCI.2826-16.2017.

(24) McGuire, S. E.; Mao, Z.; Davis, R. L. Spatiotemporal Gene Expression Targeting with the TARGET and Gene-Switch Systems in Drosophila. Sci STKE 2004, 2004 (220), pl6. 10.1126/stke.2202004pl6.

(25) Corson, F.; Couturier, L.; Rouault, H.; Mazouni, K.; Schweisguth, F. Self-Organized Notch Dynamics Generate Stereotyped Sensory Organ Patterns in Drosophila. Science 2017, 356 (6337), eaai7407. 10.1126/science.aai7407.

(26) Ganguly-Fitzgerald, I.; Donlea, J.; Shaw, P. J. Waking Experience Affects Sleep Need in Drosophila. Science 2006, 313 (5794), 1775–1781. 10.1126/science.1130408.

(27) Aso, Y.; Siwanowicz, I.; Bräcker, L.; Ito, K.; Kitamoto, T.; Tanimoto, H. Specific Dopaminergic Neurons for the Formation of Labile Aversive Memory. Curr Biol 2010, 20 (16), 1445–1451. 10.1016/j.cub.2010.06.048.

(28) Marathe, S.; Alberi, L. Notch in Memories: Points to Remember. Hippocampus 2015, 25 (12), 1481–1488. 10.1002/hipo.22426.

(29) Zhang, J.; Yin, J. C.; Wesley, C. S. From Drosophila Development to Adult: Clues to Notch Function in Long-Term Memory. Frontiers in cellular neuroscience 2013, 7, 222. 10.3389/fncel.2013.00222.

(30) Matsuno, M.; Horiuchi, J.; Tully, T.; Saitoe, M. The Drosophila Cell Adhesion Molecule Klingon Is Required for Long-Term Memory Formation and Is Regulated by Notch. Proc Natl Acad Sci U S A 2009, 106 (1), 310–315. 10.1073/pnas.0807665106.

(31) Awasaki, T.; Lai, S.-L.; Ito, K.; Lee, T. Organization and Postembryonic Development of Glial Cells in the Adult Central Brain of Drosophila. The Journal of neuroscience : the official journal of the Society for Neuroscience 2008, 28 (51), 13742–13753. 10.1523/JNEUROSCI.4844-08.2008.

(32) Kremer, M. C.; Jung, C.; Batelli, S.; Rubin, G. M.; Gaul, U. The Glia of the Adult Drosophila Nervous System. Glia 2017, 65 (4), 606–638. 10.1002/glia.23115.

(33) Miyashita, T.; Murakami, K.; Kikuchi, E.; Ofusa, K.; Mikami, K.; Endo, K.; Miyaji, T.; Moriyama, S.; Konno, K.; Muratani, H.; Moriyama, Y.; Watanabe, M.; Horiuchi, J.; Saitoe, M. Glia Transmit Negative Valence Information during Aversive Learning in Drosophila. Science 2023, 382 (6677), eadf7429. 10.1126/science.adf7429.

(34) Stork, T.; Engelen, D.; Krudewig, A.; Silies, M.; Bainton, R. J.; Klambt, C. Organization and Function of the Blood-Brain Barrier in Drosophila. The Journal of neuroscience : the official journal of the Society for Neuroscience 2008, 28 (3), 587–597. 10.1523/JNEUROSCI.4367-07.2008.

(35) Otto, N.; Marelja, Z.; Schoofs, A.; Kranenburg, H.; Bittern, J.; Yildirim, K.; Berh, D.; Bethke, M.; Thomas, S.; Rode, S.; Risse, B.; Jiang, X.; Pankratz, M.; Leimkühler, S.; Klämbt, C. The Sulfite Oxidase Shopper Controls Neuronal Activity by Regulating Glutamate Homeostasis in Drosophila Ensheathing Glia. Nat Commun 2018, 9, 3514. 10.1038/s41467-018-05645-z.

(36) Melom, J. E.; Littleton, J. T. Mutation of a NCKX Eliminates Glial Microdomain Calcium Oscillations and Enhances Seizure Susceptibility. The Journal of neuroscience : the official journal of the Society for Neuroscience 2013, 33 (3), 1169–1178. 10.1523/JNEUROSCI.3920-12.2013.

(37) Weiss, S.; Clamon, L. C.; Manoim, J. E.; Ormerod, K. G.; Parnas, M.; Littleton, J. T. Glial ER and GAP Junction Mediated Ca2+ Waves Are Crucial to Maintain Normal Brain Excitability. Glia 2022, 70 (1), 123–144. 10.1002/glia.24092.

(38) Spéder, P.; Brand, A. H. Gap Junction Proteins in the Blood-Brain Barrier Control Nutrient-Dependent Reactivation of Drosophila Neural Stem Cells. Dev Cell 2014, 30 (3), 309–321. 10.1016/j.devcel.2014.05.021.

(39) de Tredern, E.; Rabah, Y.; Pasquer, L.; Minatchy, J.; Plaçais, P.-Y.; Preat, T. Glial Glucose Fuels the Neuronal Pentose Phosphate Pathway for Long-Term Memory. Cell Rep 2021, 36 (8), 109620. 10.1016/j.celrep.2021.109620.

(40) Rabah, Y.; Francés, R.; Minatchy, J.; Guédon, L.; Desnous, C.; Plaçais, P.-Y.; Preat, T. Glycolysis-Derived Alanine from Glia Fuels Neuronal Mitochondria for Memory in Drosophila. Nat Metab 2023, 5 (11), 2002–2019. 10.1038/s42255-023-00910-y.

(41) Chaturvedi, R.; Stork, T.; Yuan, C.; Freeman, M. R.; Emery, P. Astrocytic GABA Transporter Controls Sleep by Modulating GABAergic Signaling in Drosophila Circadian Neurons. Curr Biol 2022, 32 (9), 1895–1908.e5. 10.1016/j.cub.2022.02.066.

(42) Chaturvedi, R.; Reddig, K.; Li, H.-S. Long-Distance Mechanism of Neurotransmitter Recycling Mediated by Glial Network Facilitates Visual Function in Drosophila. Proceedings of the National Academy of Sciences of the United States of America 2014, 111 (7), 2812–2817. 10.1073/pnas.1323714111.

(43) Davla, S.; Artiushin, G.; Li, Y.; Chitsaz, D.; Li, S.; Sehgal, A.; van Meyel, D. J. AANAT1 Functions in Astrocytes to Regulate Sleep Homeostasis. eLife 9, e53994. 10.7554/eLife.53994.

(44) Blum, I. D.; Keleş, M. F.; Baz, E.-S.; Han, E.; Park, K.; Luu, S.; Issa, H.; Brown, M.; Ho, M. C. W.; Tabuchi, M.; Liu, S.; Wu, M. N. Astroglial Calcium Signaling Encodes Sleep Need in Drosophila. Curr Biol 2021, 31 (1), 150–162.e7. 10.1016/j.cub.2020.10.012.

(45) Ingiosi, A. M.; Hayworth, C. R.; Harvey, D. O.; Singletary, K. G.; Rempe, M. J.; Wisor, J. P.; Frank, M. G. A Role for Astroglial Calcium in Mammalian Sleep and Sleep Regulation. Curr Biol 2020, 30 (22), 4373–4383.e7. 10.1016/j.cub.2020.08.052.

(46) Zhang, Y. V.; Ormerod, K. G.; Littleton, J. T. Astrocyte Ca2+ Influx Negatively Regulates Neuronal Activity. eNeuro 2017, 4 (2), ENEURO.0340-16.2017. 10.1523/ENEURO.0340-16.2017.

(47) Artiushin, G.; Sehgal, A. The Glial Perspective on Sleep and Circadian Rhythms. Annu Rev Neurosci 2020, 43, 119–140. 10.1146/annurev-neuro-091819-094557.

